# Investigating racial disparities in carcinomas through TCGA transcriptomic and proteomic database

**DOI:** 10.1101/2023.02.21.529336

**Authors:** Brian Lei, Xinyin Jiang, Anjana Saxena

## Abstract

**Simple Summary**: Racial disparities in cancer incidence and outcome rates are prevalent in the US, with a variety of contributing factors such as socioeconomic status, differences in lifestyle and environmental exposures, and gene polymorphisms. The goal of this research was to broadly analyze public data to identify critical differences in molecular signatures and pathways between races. Additionally, to ensure the clinical translatability of our work, we analyzed the impact of these differences on patient survival. Our findings will help inform the use of novel biomarkers in clinical settings and the future development of precision therapies.

**Abstract**:Epidemiological studies highlight a disparity in cancer incidence and outcome rates between racial groups in the United States. In our study, we investigated molecular differences among racial groups in 10 carcinoma types. We used publicly available data from The Cancer Genome Atlas to identify patterns of differential gene expression in tumors obtained from 4,112 White, Black/African American, and Asian patients. We identified race-dependent expression of numerous genes whose mRNA transcript levels were significantly correlated with patient survival. A small subset of these genes was differentially expressed in multiple carcinomas, including genes involved in cell cycle progression such as *CCNB1*, *CCNE1*, *CCNE2*, and *FOXM1*. In contrast, genes such as transcriptional factor *ETS1* and apoptotic gene *BAK1* were differentially expressed and clinically significant only in specific cancer types. Our analyses also revealed race-dependent regulation of relevant pathways. Importantly, homology directed repair and ERBB4-mediated nuclear signaling were both upregulated in Black patients compared to Whites in four carcinoma types. This large-scale pan-cancer study refines our understanding of the cancer health disparity and can help inform the use of novel biomarkers in clinical settings as well as the future development of precision therapies.

## 1. Introduction

Race is a very broad categorizer that groups populations according to common phenotypic characteristics. Racial groups are heterogenous social constructs that exhibit immense biological variation within them. Even so, the inherent differences between races are a source of great diversity in the human species and lead to underlying variation in biological mechanisms [1]. The actual extent to which race affects these mechanisms, especially those pertaining to diseases like cancer, remains to be elucidated. Cancer epidemiologists have long recognized a race-based disparity in cancer incidence and outcomes, especially in the United States. External factors, such as disparities in health insurance and access to healthcare, socioeconomic disparities, and differences in risk factor exposure (e.g., pollution) play a substantial role as well [2]. For example, American cancer registries have shown that cancer survival rates are lower in Blacks at each stage of diagnosis across a wide spectrum of cancer types [3]. Some studies have suggested that differences in outcome persist even under equal-access care environments [4], while others suggest that no differences persist in cancer-specific mortality rates after adjustment for non-biological factors [5]. Such results are complicated by further stratifications along the lines of sex, age, and cancer type. For example, analyses of longitudinal National Cancer Institute Surveillance, Epidemiology, and End Results (SEER) program data conducted by Zeng et al. found significantly improved survival in younger patients for a variety of major cancer types but no apparent sex difference. [6] On the other hand, another analysis of SEER data done by Dong et al. found significantly higher incidence and worse survival outcomes in men compared to women in the majority of studied cancer types. [7]

Recent research provides extensive insight into the disparity as it occurs in various cancer types. For example, prostate cancer occurs more often and has greater mortality rates in African American men as compared to Caucasian Americans, which has been increasingly linked to genetic and molecular alterations in addition to socioeconomic factors [8]. Likewise, incidence rates of aggressive endometrial cancers are significantly higher in non-Hispanic Black women compared to White women, and 5-year relative survival for Black women is significantly less than Whites [9]. Nipp et al. found that in early-stage pancreatic adenocarcinoma, Black and Hispanic patients suffer from worse survival than whites [10]. Islami et al. identified a major ethnic disparity in liver cancer death rates linked to differences in risk factor prevalence, ranging from 5.5 per 100,000 in non-Hispanic Whites to 11.9 per 100,000 in American Indians/Alaska Natives [11]. A racial examination of the California Cancer Registry revealed that advanced stage and high-grade bladder cancer is especially prevalent in Black patients, along with significantly poorer 5-year disease-specific survival [12]. Tannenbaum et al. identified improved survival for non-small cell lung cancer in Asians compared to Whites in Florida [13]. On the other hand, Tramontano et al. found that there was no significant disparity of mortality between White and Nonwhite esophageal cancer patients after adjustment [14].

These racial disparities are linked to a number of differentially expressed genes (DEGs). For example, Li et al. identified the *XKR9* gene, implicated in the exposure of phosphatidylserine during apoptosis, as being differentially expressed between Asian Americans, Caucasian Americans, and African Americans in many cancer types and significantly associated with overall survival [15]. Thus, *XKR9* could act as a potential race-dependent target for cancer immunotherapy. However, the results of smaller-scale studies suggest that patterns of differential expression vary significantly across cancer types: Grunda et al. identified differential expression of the prognostically significant genes *AR, BCL2, CCND1, CDKN1A, CDKN1B, CDKN2A, ERBB2, ESR1, GATA3, IGFBP2, IL6ST, KRT19, MUC1, PGR*, and *SERPINE1* in non-Hispanic White and African American breast cancer patients [16]. Vazquez et al. used TCGA data to discover differential expression of chemokine receptors depending on race and molecular subtype, which could explain racial differences in tumor microenvironment and immunotherapy response [17]. Powell et al. highlighted race-dependent expression of genes such as *AKT1, ALOX12, IL8, CXCR4, FASN*, and *TIMP3* in prostate cancer, suggesting opportunities for targeted therapies [18]. Other prostate cancer research points to aberrant activation of pathways such as androgen receptor (AR) signaling, epidermal growth factor receptor (EGFR) signaling, and inflammatory signaling across races as potential causes of the disparity [19]. **Table 1** depicts many such genes that are categorized as differentially expressed genes (DEGs) in cancer tissue in Whites vs. Blacks that encompass a variety of cancer types and influence many critical biological pathways. Additionally, other biological factors, such as genetic polymorphism, mutational variation, and epigenetic variation have also been linked to race-based differences in cancer incidence and outcome rates. For example, Asif et al. discovered increased CpG hypomethylation in Black endometrial cancer patients compared to Whites, indicating worsened oncogenic transformation [20].

**Table 1.** Differentially expressed genes in breast, colorectal, endometrial, lung, and prostate cancer, and their relevant pathways.

| Cancer type | Higher expression | DEGs | References |
| --- | --- | --- | --- |
| **All** | Black | IGF1, IL2RA, IL6 | [1,15] |
|  | White | XKR9, CST1, MTRNR2L1 |  |
| **Breast** | Black | CDKN2A, CRYBB2, SRC, SCN1A | [1,16,21] |
|  | White | AR, BCL2, CASP8, CCND1, CDKN1A, CDKN1B, ERBB2, ESR1, GATA3, IGFBP2, IL6ST, KRT19, MUC1, PGR, SERPINE1 |  |
| **Colorectal** | Black | EGFR, TOP2A, CRYBB2, PSPH, IL33 | [22,23] |
|  | White | IL27, ADAL, ARSE, SMOC1, HESX1, TRNP1 |  |
| **Endometrial** | Black | MCM2, PLK1, MCM7, LAMA5, LAMC1 | [24] |
|  | White | SMC1B, CDC7, CCNE2, BRCA2, ERBB2, FGFR3 |  |
| **Lung** | White | CD274 | [22] |
| **Prostate** | Black | AKT1, SPINK1, AR, ARA55, GNB3, POLR2L, TBP, EGFR, CRYBB2, IL6, IL8, CXCR4, FASN | [1,18,19] |
|  | White | ALOX12, TIMP3 |  |

In this study we aimed to analyze pre-existing datasets from The Cancer Genome Atlas (TCGA) with a focus on molecular differences between White, African American/Black, and Asian patients. While poorer cancer prognoses and incidence rates have been associated with specific racial groups [3,25], the molecular mechanisms responsible for this disparity remain the subject of ongoing research. Large genomic, transcriptomic and proteomic datasets available from a variety of studies and patient groups are often under-analyzed with regard to racial analysis. In this study, we assessed these large datasets with the lens of race to gain insights about the racial dependency of DEGs, biological pathways, and patient survival. These revelations will be pivotal not only in designing and developing precision therapies in the future, but also in providing cancer-specific sets of biomarkers that can be useful in clinical diagnoses and prognoses.

## 2. Methodology

### 2.1. Data information

We used TCGA datasets generated as part of the PanCancer Atlas tumor molecular analysis project, catalogued in **Table 2**. These PanCancer Atlas datasets are already normalized to facilitate comparative analyses. We accessed these open-access, open-source cancer database and analyzed using web analysis tool cBioPortal [26–28]. The selected studies encompassed ten different cancer types, including the PanCancer datasets of colorectal adenocarcinoma (CORE, n = 338), uterine corpus endometrial carcinoma (UCEC, n = 445), invasive breast carcinoma (BRCA-P, n = 903), kidney renal clear cell carcinoma (KIRC, n = 383), kidney renal papillary cell carcinoma (KIRP, n = 253), liver hepatocellular carcinoma (LIHC, n = 329), lung adenocarcinoma (LUAD, n = 434), stomach adenocarcinoma (STAD, n = 344), and thyroid carcinoma (THCA, n = 370) [29]. In addition to these nine PanCancer Atlas datasets, we also included and analyzed two TCGA datasets profiled in 2015 *Cell* studies, namely Invasive Breast Carcinoma (BRCA-C, n = 684) [30] and Prostate Adenocarcinoma (PRAD, n = 313) [31]. These two datasets also included gene methylation data. PRAD was the only study that did not include survival information associated with its samples. For the purposes of race-based survival, mRNA, protein, and methylation analysis, all data were filtered in the cBioPortal web tool to only include complete samples, or samples that included associated mRNA expression data, copy number alteration data, and mutation data.

**Table 2.**
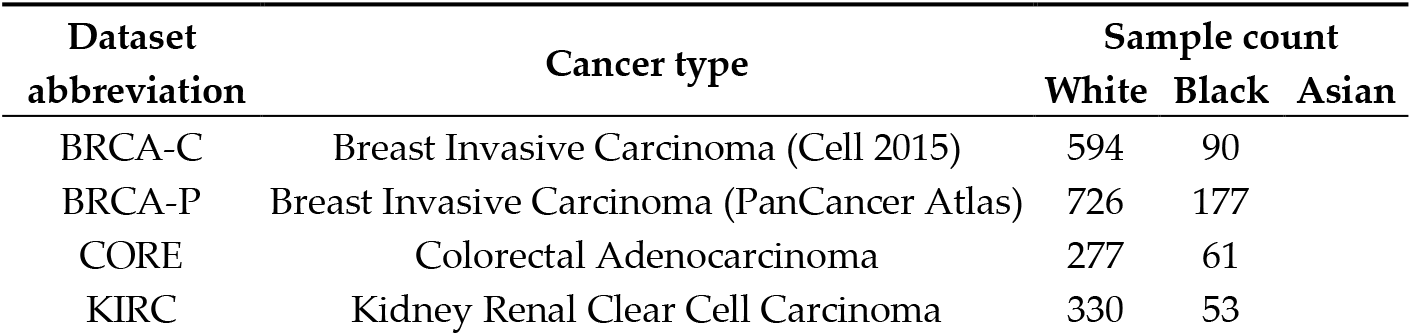
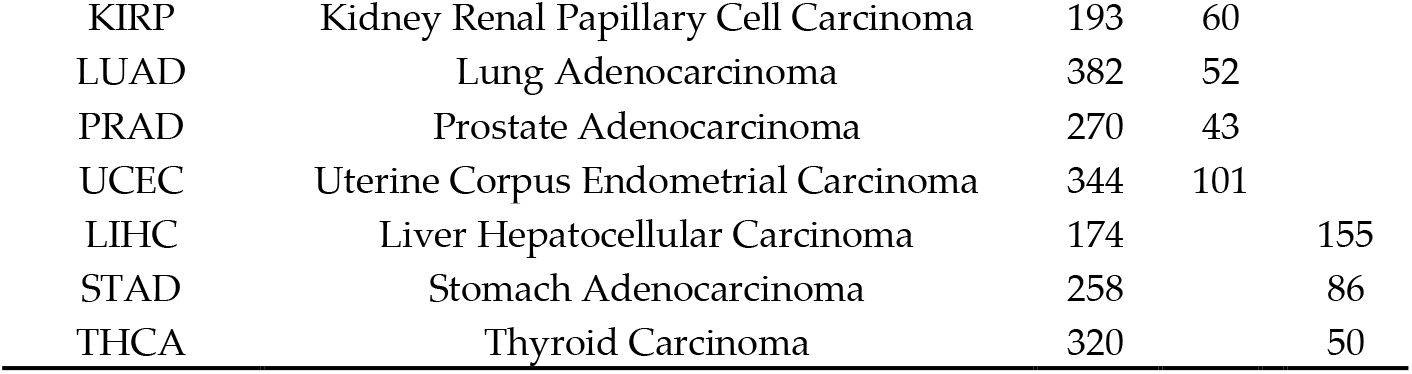
Breakdown of selected cancer datasets and respective sample counts. One sample corresponds to one patient. Gene expression analyses were performed between two races at a time for a given cancer type: Whites vs. Blacks/African Americans or Whites vs. Asians.

### 2.2. Race-based Kaplan-Meier survival analysis

We performed race-based survival analysis for each cancer type in the cBioPortal web tool using race as the comparison factor. For BRCA-C, BRCA-P, CORE, KIRC, KIRP, LUAD, and UCEC, we compared “White” labeled samples against “Black/African American” labeled samples. For LIHC, STAD, and THCA, samples labeled “White” were compared against samples labeled “Asian”. In addition to this baseline racial analysis, we performed separate analyses after stratifying the racial groups by sex (e.g., White males vs. Black males, White females vs. Black females). The particular racial groups we selected for analysis varied due to unequal levels of racial representation in the different cancer datasets. Survival was evaluated using four different survival metrics: disease-free survival, progression-free survival, disease-specific survival, and overall survival. We used log rank tests performed by cBioPortal to assess statistical significance; p ≤ 0.05 was used as the threshold of significance.

### 2.3. Gene-based Kaplan-Meier survival analysis

To determine the most clinically relevant genes for each cancer type, we used the Kaplan-Meier (KM) Plotter web tool [32]. The Kaplan-Meier plotter database integrates expression data and clinical outcomes from the GEO, EGA, and TCGA [33]. We evaluated the prognostic value of the mRNA expression levels of a compiled set of 155 target oncogenes and tumor suppressor genes (see **S1**) for all cancer types listed in **Table 2**, except for prostate adenocarcinoma for which data were unavailable and hence excluded from this analysis.

For each gene, patients were divided into two groups of low and high mRNA expression based on the median expression and the overall survival rates were compared. P-values and hazard ratios with 95% confidence intervals were calculated. To reduce the false discovery rate, we used p ≤ 0.01 as a threshold of significance.

### 2.4. mRNA, protein, and methylation analysis

In the cBioPortal web tool, we performed group comparison analyses with race as the comparison factor. For BRCA-C, BRCA-P, CORE, KIRC, KIRP, LUAD, PRAD, and UCEC, we compared “White” labeled samples against “Black/African American” labeled samples. For LIHC, STAD, and THCA, samples labeled “White” were compared against samples labeled “Asian”. The racial groups selected for analysis varied due to unequal levels of racial representation in the different cancer datasets. For each study, mRNA and protein expression analysis results were downloaded as TSV files, as well as DNA methylation analysis results for BRCA-C and PRAD. We wrote and used a pandas-based Python (version 3.7.7) program to filter the data for the 155 target genes. Statistically significant differences in mRNA and protein expression between the two racial cohorts, as determined by Benjamini-Hochberg procedure performed by the cBioPortal web tool, were identified.

### 2.5. Reactome pathway analysis

To perform integrative pathway analysis, we downloaded the original PanCancer Atlas RPPA microarray and clinical data from the PanCancer Atlas publication page [34,35]. This data was not filtered for the 155 target genes for the purposes of pathway analysis. We created a pandas-based Python program (https://github.com/brian-lei/cancer-racial-disparity) to format and export the data to Reactome, an open-source, open access peer-reviewed pathway database [36,37]. We grouped samples by cancer type as well as by race, deleted duplicate samples, and removed protein data for a select few proteins that contained many corresponding missing values, namely ADAR1, alpha-catenin, TTF-1, caspase-3, caspase-9, PARP1, and JAB1. Post-translationally modified versions of proteins were also excluded from the analysis. We manually annotated samples according to their corresponding race in the Reactome web tool and performed a Pathway Analysis with Down-weighting of Overlapping Genes (PADOG) microarray analysis with race as the comparison factor.

### 2.6. Statistical analysis as presented in Results

Genes that received p-values less than or equal to 0.01 in gene-based Kaplan-Meier (KM) survival analysis were considered “clinically significant”. The survival relationship, as evaluated by KM analysis, is denoted in tables with a “+” for positive correlation between gene mRNA expression and overall survival and a “–“ for negative correlation between gene mRNA expression and overall survival, followed by the associated log rank p-value in parentheses.

Genes that received q-values less than or equal to 0.05 in cBioPortal Ben-jamini-Hochberg comparison analysis for mRNA expression, protein expression, or both were considered to be “differentially expressed”. The race denoted in tables under “mRNA” is the racial group with higher mRNA expression of the gene, followed by the associated q-value in parentheses. The race denoted in tables under “Protein” is the race with higher protein expression and the associated q-value. “NS” represents non-significant values. Empty cells represent missing data from the original TCGA database.

Each table in the results section presents gene expression information for a particular cancer type. In all cases, we only included genes that were both clinically significant and differentially expressed between the denoted races. We excluded genes for which neither mRNA nor protein expression was significantly different.

In the Reactome Pathway Analysis subsection, pathways found differentially expressed at an adjusted p-value less than or equal to 0.05 were considered significant. The adjusted p-value is denoted in tables in parentheses in each table header.

## 3. Results

The analysis results are organized alphabetically by cancer type. Each section reports the results of cBioPortal differential expression analysis. All sections except for Prostate Adenocarcinoma include cBioPortal race-based survival and KM gene-based survival analysis results. Gene methylation analysis was performed for only two studies for which data were available: Breast Invasive Carcinoma (BRCA-C) and Prostate Adenocarcinoma (PRAD). Gene expression analysis was performed on only one dataset for each section/cancer type, except for Breast Invasive Carcinoma which includes results from both BRCA-C and BRCA-P. Complete results on all 155 target genes are included in **S2-S4**. The original Reactome report is included in **S5**.

### 3.1. Breast Invasive Carcinoma

#### 3.1.1. Race-based survival

In BRCA-C and BRCA-P, no significant differences were identified between Whites and Blacks/African Americans for any of the survival metrics.

#### 3.1.2. Gene-based survival

High BCL3, FHIT, and IGF1R transcript levels were correlated with increased overall patient survival, while high CCNE1 and TGFBR1 transcript levels were correlated with decreased overall patient survival.

#### 3.1.3 Differential expression

BRCA-C and BRCA-P data revealed a relatively high number of differentially expressed genes compared to the other studies. Most notably, in BRCA-C, CCNE1 mRNA and protein levels were significantly increased in Black patients compared to White patients. However, in BRCA-P, protein levels were not significantly different (**Table 3**).

**Table 3.**
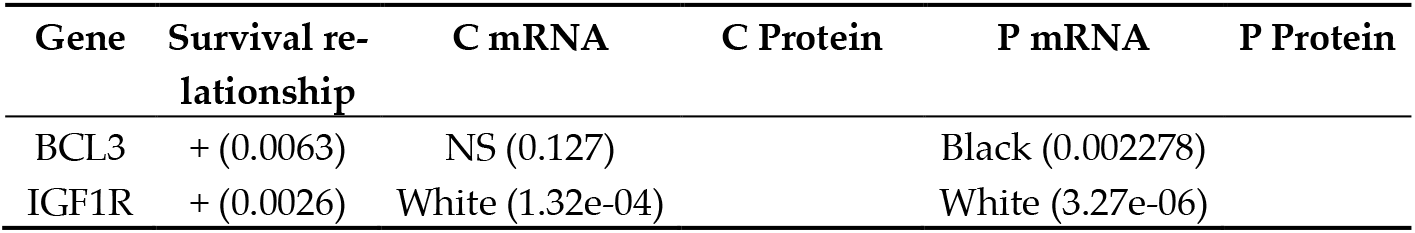
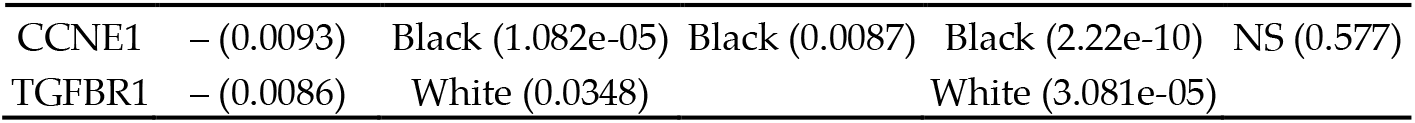
BRCA-C and BRCA-P (indicated as C and P respectively), breast invasive carcinoma. Out of all 155 target genes, five genes were identified by KM analysis as clinically significant. Of these, the four shown genes were differentially expressed between Black and White patients in at least one of the two datasets.

87 other non-clinically significant genes exhibited differential expression across races. For example, BAK1, CCNB1, NOTCH1, RAD51, and STMN1 mRNA and protein levels were enriched in Black patients in both studies, while KDR, MAPK9, and RAD50 mRNA and protein were enriched in White patients in both studies. Many other genes, such as APC, AXL, CDK4, CXCR1, ERCC4, IGF1, IGF1R, KRAS, MYB, PDGFD, RASSF8, and TGFBR1 had similar patterns of differential mRNA expression in both studies, but lacked corresponding protein data.

#### 3.1.4. Methylation

In BRCA-C, CCNE1 methylation was significantly increased in White patients compared to Black patients. 38 other target genes exhibited differential methylation across racial groups, including BRCA2 (higher methylation in Whites) and PTEN (higher methylation in Blacks).

### 3.2. Colorectal Adenocarcinoma

#### 3.2.1. Race-based survival

In CORE, Blacks and African Americans suffered from significantly worse progression-free survival (p = 0.0476) than Whites, while disparities in disease-free (p = 0.443), disease-specific (p = 0.0513), and overall survival (p = 0.961) were not significant. While survival of Black females and White females did not differ significantly in any metric, Black males had significantly worse progression-free (p = 0.004526) and disease-specific survival (p = 0.0362) than White males.

#### 3.2.2. Gene-based survival

Out of all 155 target genes, only BIRC2 and FGF16 transcript levels had a significant impact on survival in rectal adenocarcinoma. High BIRC2 transcript levels were correlated with increased overall survival, while high FGF16 transcript levels were correlated with decreased survival.

#### 3.2.3. Differential expression

While BIRC2 mRNA expression was not significantly different between Whites and Blacks/African Americans, BIRC2 protein expression was significantly increased in Blacks (**Table 4**). Unfortunately, neither mRNA nor protein data were available for FGF16 in the original TCGA data despite being clinically significant.

**Table 4.**
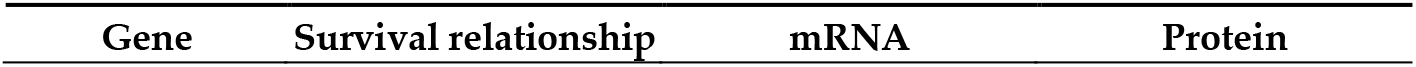
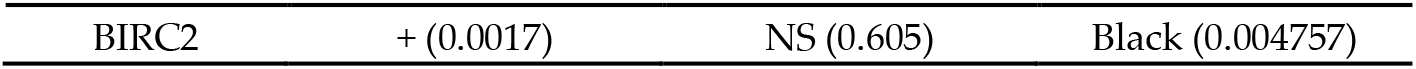
CORE, colorectal adenocarcinoma. Out of all 155 target genes, two (BIRC2 and FGF16) were identified by KM analysis as clinically significant. Of these, only BIRC2 was differentially expressed between White and Black/African American patients.

For the other 153 non-clinically significant genes, except for DIABLO and FHIT, the transcript levels either did not differ significantly between Black and White patients or did not exist in the original TCGA data. DIABLO mRNA expression was higher in Whites (protein levels were not significantly different) while FHIT expression was higher in Blacks (protein data not available). 24 genes exhibited differential protein expression: BAK1, BIRC2, FASN, FOXM1, IRS1, MSH2, and STMN1 protein levels were significantly increased in Black patients while CCNE2, CDH2, CDKN1A, CHEK1, DIRAS3, ERCC1, ETS1, GAPDH, HSPA1A, MAPK9, MYC, NF2, PRDX1, TGM2, TIGAR, VHL, and XRCC1 protein levels were significantly increased in White patients.

### 3.3. Kidney Renal Clear Cell Carcinoma

#### 3.3.1. Race-based survival

No significant differences were identified between Whites and Blacks/African Americans for any of the survival metrics.

#### 3.3.2. Gene-based survival

High APC, BCL2, CCND1, CDKN1B, CTNNB1, ERBB2, ERBB3, ERCC4, ERG, ETS1, FGF1, FGF12, FLT1, GAB2, IGF1R, IL6R, KDR, KRAS, MAPK1, MAPK9, MSH2, MYCN, NRAS, PDGFC, PDGFD, PTCH1, PTEN, RAD50, RASSF8, RB1, SCFD2, SMAD4, TAL2, TGFA, TGFBR2, and TGFBR3 transcript levels were correlated with increased overall patient survival.

High AXL, BAK1, BCL3, CCNB1, CCNE1, CCNE2, CRP, ERCC1, ETV4, FASN, FGF5, FGF8, FGF17, FGF21, FGF23, FOXM1, GAPDH, GNAS, GRP, IL6, NKX2-5, NKX3-1, NTRK1, SHBG, and SRC transcript levels were correlated with decreased overall patient survival.

#### 3.3.3. Differential expression

AXL, FGF5, KRAS, NKX2-5, RB1, and TAL2 transcript levels were significantly higher in White patients compared to Black patients although protein data were absent from the original TCGA data. NRAS transcript levels were also increased in White patients but protein expression was not significantly different across races. ETV4 and FGF17 transcript levels were higher in Black patients although protein data were absent. ERBB2 transcript levels were higher in Black patients but protein did not differ significantly across races. ETS1, MAPK1, MAPK9, and RAD50 mRNA and protein levels were both significantly increased in White patients. BCL2, CCNE2, ERCC1, and GAPDH protein levels were also increased in Whites, although transcript levels were not significantly different across races. Interestingly, CCNB1, MSH2, FOXM1, and RAD51 protein levels were increased in Black patients while transcript levels were increased in White patients (**Table 5**).

**Table 5.**
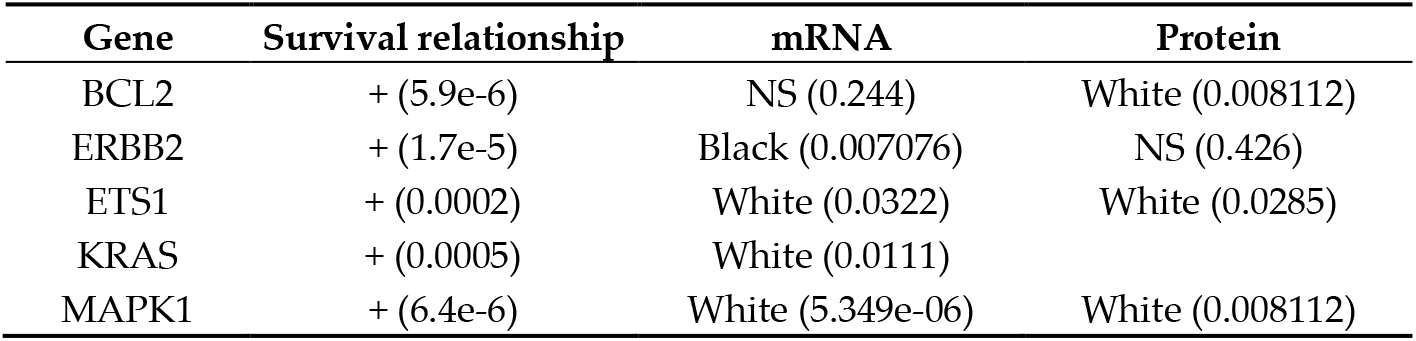
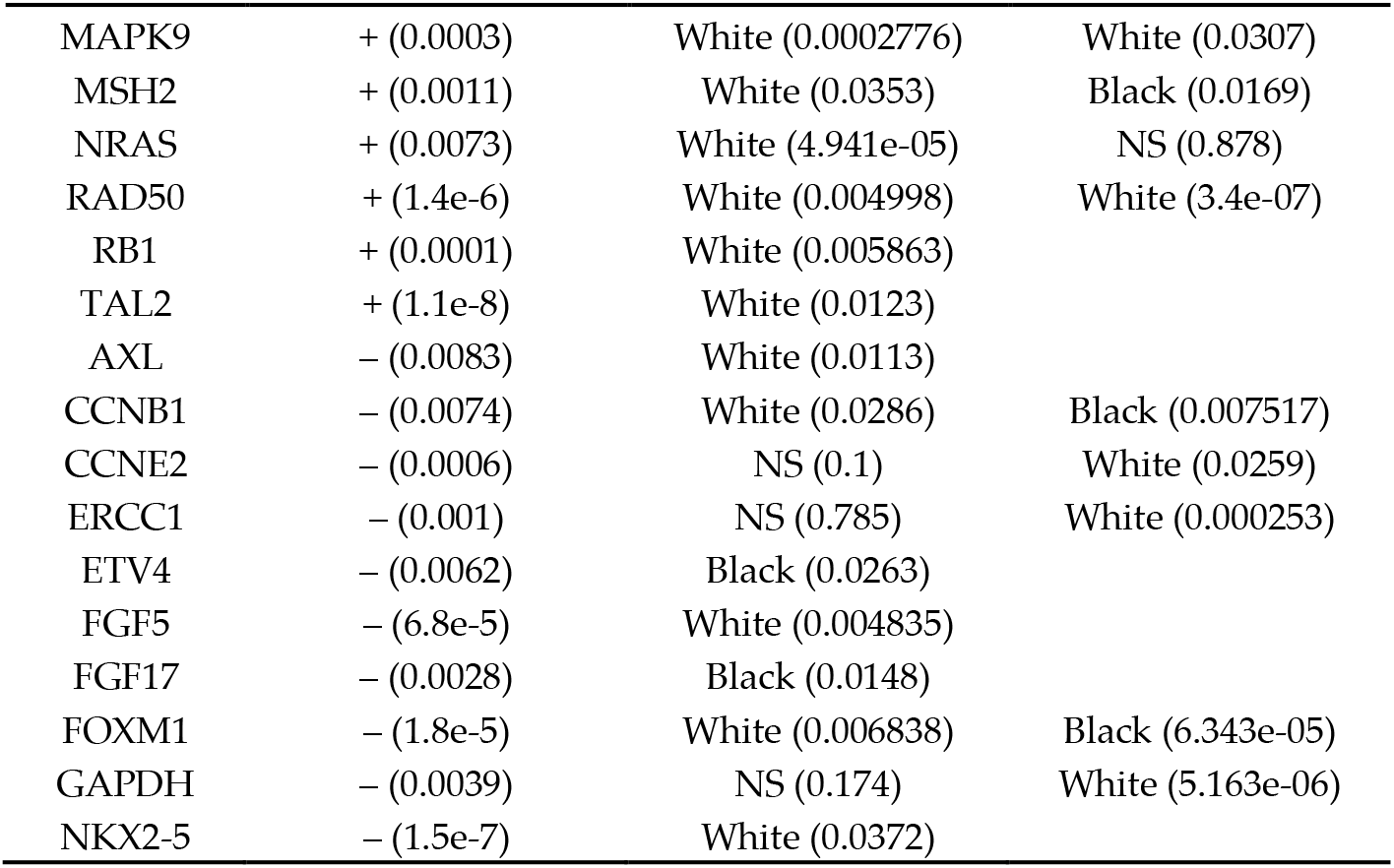
KIRC, kidney renal clear cell carcinoma. Out of all 155 target genes, 61 genes were clinically significant in KM analysis; 21 genes were differentially expressed between Black and White patients as depicted below.

| Gene | Survival relationship | mRNA | Protein |
| --- | --- | --- | --- |
| BCL2 | + (5.9e-6) | NS (0.244) | White (0.008112) |
| ERBB2 | + (1.7e-5) | Black (0.007076) | NS (0.426) |
| ETS1 | + (0.0002) | White (0.0322) | White (0.0285) |
| KRAS | + (0.0005) | White (0.0111) |  |
| MAPK1 | + (6.4e-6) | White (5.349e-06) | White (0.008112) |
| MAPK9 | + (0.0003) | White (0.0002776) | White (0.0307) |
| MSH2 | + (0.0011) | White (0.0353) | Black (0.0169) |
| NRAS | + (0.0073) | White (4.941e-05) | NS (0.878) |
| RAD50 | + (1.4e-6) | White (0.004998) | White (3.4e-07) |
| RB1 | + (0.0001) | White (0.005863) |  |
| TAL2 | + (1.1e-8) | White (0.0123) |  |
| AXL | – (0.0083) | White (0.0113) |  |
| CCNB1 | – (0.0074) | White (0.0286) | Black (0.007517) |
| CCNE2 | – (0.0006) | NS (0.1) | White (0.0259) |
| ERCC1 | – (0.001) | NS (0.785) | White (0.000253) |
| ETV4 | – (0.0062) | Black (0.0263) |  |
| FGF5 | – (6.8e-5) | White (0.004835) |  |
| FGF17 | – (0.0028) | Black (0.0148) |  |
| FOXM1 | – (1.8e-5) | White (0.006838) | Black (6.343e-05) |
| GAPDH | – (0.0039) | NS (0.174) | White (5.163e-06) |
| NKX2-5 | – (1.5e-7) | White (0.0372) |  |

37 non-clinically significant genes exhibited differential expression across races. Interestingly, BAP1 transcript levels were increased in Black patients while protein levels were higher in White patients. Several other genes had differential protein expression but not differential mRNA expression, namely BECN1, BIRC2, CDKN1A, DIABLO, DIRAS3, IRS1, NF2, PRDX1, RAF1, STMN1, TGM2, TP53, VHL, and XRCC1.

### 3.4. Kidney Renal Papillary Cell Carcinoma

#### 3.4.1. Race-based survival

No significant differences were identified between Whites and Blacks/African Americans for any of the survival metrics.

#### 3.4.2. Gene-based survival

High AXL and SMAD4 transcript levels were correlated with increased overall patient survival, while high BRCA2, CCNE1, CCNE2, FASN, FGF5, FGF7, FGF8, FGF11, FGF18, FOXM1, GRP, IGF2, and PDGFRB transcript levels were correlated with decreased overall patient survival.

#### 3.4.3. Differential expression

For all 155 target genes, including the genes identified by KM analysis as clinically significant for kidney renal papillary cell carcinoma, mRNA expression was not significantly different across races. Differences in protein expression were either not significant (e.g., BRCA2) or indeterminable due to a lack of associated protein data (e.g., AXL).

### 3.5. Liver Hepatocellular Carcinoma

#### 3.5.1. Race-based survival

No significant differences were identified between Whites and Asians for any of the survival metrics.

#### 3.5.2. Gene-based survival

High BAK1, CCNB1, CCNE2, CDK4, CDKN2A, CHEK1, DIABLO, ETV1, ETV4, FOXM1, GAPDH, HRAS, MEN1, MSH2, NKX2-5, NRAS, PRDX1, RAD51, RASSF7, STMN1, TIGAR, and VHL transcript levels were all correlated with decreased overall patient survival. Interestingly, none of the target genes were correlated with improved survival.

#### 3.5.3. Differential expression

In LIHC, protein data for the genes identified by KM analysis as clinically relevant for liver hepatocellular carcinoma were either absent or displayed non-significant differences across races, except for BAK1 whose protein levels were significantly increased in White patients compared to Asian patients. BAK1 transcript levels, however, were not significantly different across races. CCNB1, CCNE2, CHEK1, FOXM1, HRAS, MSH2, RAD51, and STMN1 transcript levels were significantly higher in Asian patients, while TIGAR transcripts were increased in White patients (**Table 6**).

**Table 6.** LIHC, liver hepatocellular carcinoma. Out of all 155 target genes, 22 were identified by KM analysis as clinically significant. Of these, the following 10 were differentially expressed between White and Asian patients. Interestingly, all 10 genes were negatively correlated with patient overall survival.

| Gene | Survival relationship | mRNA | Protein |
| --- | --- | --- | --- |
| BAK1 | – (0.008) | NS (0.366) | White (0.0446) |
| CCNB1 | – (0.0008) | Asian (0.004704) | NS (0.176) |
| CCNE2 | – (0.0011) | Asian (0.002332) | NS (0.22) |
| CHEK1 | – (0.0002) | Asian (0.0256) | NS (0.22) |
| FOXM1 | – (0.0015) | Asian (0.0238) | NS (0.374) |
| HRAS | – (0.0012) | Asian (0.0375) |  |
| MSH2 | – (0.0004) | Asian (0.00474) | NS (0.152) |
| RAD51 | – (0.007) | Asian (0.003367) | NS (0.302) |
| STMN1 | – (0.0004) | Asian (0.01) | NS (0.731) |
| TIGAR | – (0.0092) | White (0.002248) | NS (0.857) |

31 non-clinically significant genes exhibited differential expression across races; all of these genes differed in mRNA expression only. Specifically, AKT1, ALK, BCL2, CDKN1A, CRP, EGFR, ERCC4, ETS1, FGF7, FLT1, GDNF, GRP, HBEGF, IGF1, IL6, IL6R, KIT, MDM2, NKX3-1, PDGFC, PDGFD, PTCH1, PTCH2, RASSF6, RET, TGFBR1, TGFBR2, and TGFBRAP1 mRNA levels were enriched in White patients, while CCNE1, GNAS, and MAPK9 mRNA levels were enriched in Asian patients. Protein expression data for these genes were either not significant or unavailable from the original TCGA database.

### 3.6. Lung Adenocarcinoma

#### 3.6.1. Race-based survival

No significant differences were identified between Whites and Blacks/African Americans for any of the survival metrics.

#### 3.6.2. Gene-based survival

High BCL2, FGF18, GAB2, KIT, NTRK1, PTCH1, RASSF2, and TMPRSS2

transcript levels were correlated with increased overall patient survival, while high CCNB1, CCNE2, FGF5, FGF19, FOXM1, GAPDH, IRS1, TGFA, and VEGFC transcript levels were correlated with decreased survival.

#### 3.6.3. Differential expression

mRNA transcript levels were not significantly different between White and Black/African American patients for any of the genes identified by KM analysis as clinically significant for lung adenocarcinoma. Notably, IRS1 protein levels were significantly increased in Black patients (**Table 7**).

**Table 7.** LUAD, lung adenocarcinoma. Out of all 155 target genes, 17 were identified by KM analysis as clinically significant. Of these, only IRS1 exhibited differential expression.

| Gene | Survival relationship | mRNA | Protein |
| --- | --- | --- | --- |
| IRS1 | – (0.0099) | NS (0.669) | Black (0.0486) |

Seven non-clinically significant genes exhibited differential expression across races. ERCC1 and TIGAR protein expression was significantly increased in White patients, while NOTCH1 and RAD51 protein levels were increased in Black patients. mRNA levels for these four genes were not different between Whites and Blacks. MAPK1, MAPK9, and TGFBR1 transcript levels were enriched in Whites compared to Blacks; protein data were not significant for MAPK1/MAPK9 and absent from the original data for TGFBR1.

### 3.7. Prostate Adenocarcinoma

#### 3.7.1. Differential expression

mRNA transcript levels of CTNNB1, FGF19, NF2, PTCH1, and RB1 were elevated in White patients while transcript levels of FGFR4 and RASSF7 were elevated in Black patients. Protein data for these genes were either not significant or missing. SMAD4 protein levels were increased in White patients, although mRNA levels were not significantly different between Blacks and Whites.

#### 3.7.2. Methylation

No significant methylation differences were identified in PRAD for any of the 155 target genes.

### 3.8. Stomach Adenocarcinoma

#### 3.8.1 Race-based survival

In STAD, Asians suffered from significantly worse disease-free survival (p = 0.0145) than Whites, while disparities in progression-free (p = 0.729), disease-specific (p = 0.901), and overall survival (p = 0.304) were not significant. White females suffered from significantly worse overall survival compared to Asian females (p = 0.0453), while Asian males suffered from significantly worse disease-free survival compared to White males (p = 0.009962).

#### 3.8.2. Gene-based survival

High MSH2 transcript levels were correlated with increased overall patient survival, while high EGF, GRP, PDGFD, PDGFRB, TGFBR1, and VEGFC transcript levels were correlated with decreased overall patient survival.

#### 3.8.3. Differential expression

mRNA transcript levels were not significantly different between White and Asian patients for any of the genes identified by KM analysis as clinically significant for stomach adenocarcinoma. The exception is GRP, whose transcript data were absent from the original database. Protein data were absent for all of the clinically important genes with the exception of MSH2 protein, which was not significantly different between White and Asian patients.

Two non-clinically significant genes, PRDX1 and VHL, exhibited differential expression across races. While PRDX1 transcript levels were not significantly different, protein levels were increased in White patients. VHL mRNA expression was increased in Asian patients while protein levels were not different.

### 3.9. Thyroid Carcinoma

#### 3.9.1. Race-based survival

No significant differences were identified between White and Asian patients for any of the survival metrics.

#### 3.9.2. Gene-based survival

High EGF and FGF5 transcript levels were correlated with decreased overall patient survival.

#### 3.9.3. Differential expression

Out of the 155 target genes, none exhibited differential protein expression between White patients and Asian patients. EGF and FGF5 mRNA expression did not differ significantly between Whites and Asians, and protein data for both genes were absent from THCA. The sole significant result was that TIGAR, a non-clinically significant gene, had enriched transcript levels in Whites.

### 3.10. Uterine Corpus Endometrial Carcinoma

#### 3.10.1. Race-based survival

In UCEC, Blacks and African Americans suffered from significantly worse disease-specific survival (p = 0.0392) while disparities in progression-free (p = 0.263), disease-free (p = 0.272), and overall survival (p = 0.331) were not significant.

#### 3.10.2. Gene-based survival

High FGF1, IGF1, and MDM2 transcript levels were correlated with increased overall patient survival, while high ALK, BAK1, CCNE1, CDKN2A, FGF11, FGF12, MSH2, and RET transcript levels were correlated with decreased overall patient survival.

#### 3.10.3. Differential expression

MDM2 transcript levels were significantly higher in White patients while FGF12 and CDKN2A transcripts were higher in Black patients; protein data were absent for these genes in the original TCGA data. CCNE1 mRNA and protein levels were both increased in Black patients compared to White patients. Additionally, while MSH2 mRNA levels were not significantly different across races, protein levels were elevated in Black patients (**Table 8**).

**Table 8.** UCEC, uterine corpus endometrial carcinoma. Out of all 155 target genes, 11 were identified by KM analysis as clinically significant. Of these, the following 5 exhibited differential expression between White and Black/African American patients.

| Gene | Survival relationship | mRNA | Protein |
| --- | --- | --- | --- |
| MDM2 | + (0.0013) | White (0.016) |  |
| CCNE1 | − (0.0001) | Black (0.0298) | Black (0.0146) |
| CDKN2A | − (9.7e-5) | Black (0.0104) |  |
| FGF12 | − (0.0042) | Black (0.00292) |  |
| MSH2 | − (0.0081) | NS (0.106) | Black (0.0277) |

30 non-clinically significant genes exhibited differential expression across races. Specifically, ABL1, ETV1, FGF2, FGF5, GLI1, PDGFRA, PDGFRB, RASSF6, and TP53 transcript levels were elevated in White patients while CASP12, CDH2, MYCL, and VEGFD transcript levels were elevated in Blacks. Protein data for these genes were either not significant or absent from the original data.

Additionally, BECN1, BIRC2, ERBB3, FASN, FOXM1, IRS1, NRAS, RAD51, and STMN1 protein levels were significantly increased in Black patients while CDKN1A, CTNNB1, ERCC1, ETS1, GAPDH, RAD50, and TGM2 protein levels were increased in White patients. However, mRNA levels were not significantly different across races for any of these genes. Interestingly, while NOTCH1 mRNA expression was significantly higher in White patients, protein levels were enriched in Black patients.

### 3.11. Reactome Pathway Analysis

Homology-directed repair (HDR) was differentially regulated between Black and White patients in BRCA, CORE, KIRC, LUAD, and UCEC due to the differential protein expression of ten genes, namely ATM, BRCA2, CHEK1, ERCC1, MRE11, PCNA, RAD50, RAD51, TP53BP1, and XRCC1 (**Table 9**). In CORE, HDR was overall downregulated in Blacks compared to Whites (adjusted p = 0.027), while in the four other cancer types, HDR was upregulated in Black patients.

**Table 9.**
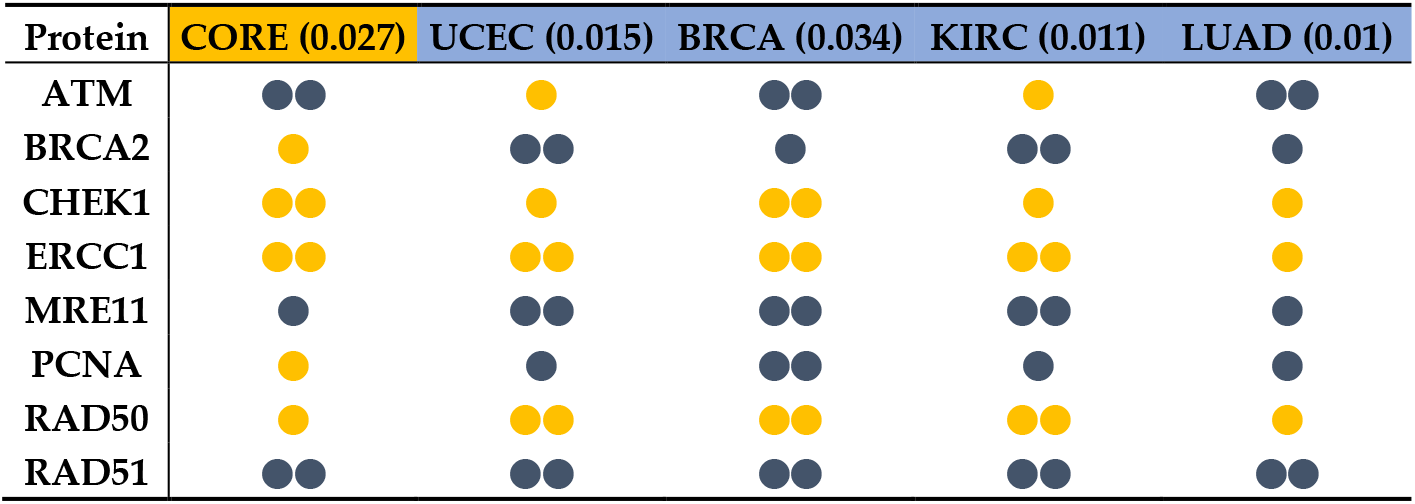
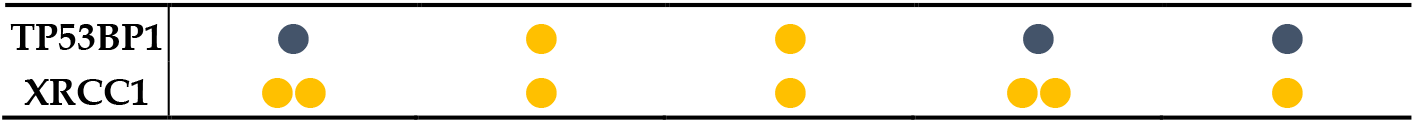
Differential protein expression for homology-directed repair genes in CORE, UCEC, BRCA, KIRC, and LUAD, compared for Black/African Americans vs. Whites. Two blue or yellow dots indicate significantly higher protein expression in Blacks or Whites, respectively. A color-coded single dot represents non-significantly higher expression in the corresponding race.

In CORE, nucleotide excision repair was significantly downregulated in Black patients compared to White patients (adjusted p = 0.019) due to decreased XRCC1 and ERCC1 protein expression.

In LIHC, DNA mismatch repair pathways were upregulated in Asian patients compared to White patients due to elevated MSH6 protein levels (adjusted p = 0.024). Additionally, although the G1/S DNA damage check-point pathway was not differentially regulated in any of the datasets, the G2/M DNA damage checkpoint pathway was upregulated in Asian patients in LIHC (adjusted p = 0.018). In Asians, ATM levels were higher while CDK1, TP53, YWHAB, and YWHAE levels were lower compared to Whites.

In BRCA, CORE, KIRC, and LUAD, ERBB4-mediated nuclear signaling was upregulated in Black patients compared to White patients due to the differential protein expression of six genes, namely ESR1, PGR, SRC, STAT5A, STMN1, and YAP1 (**Table 10**).

**Table 10.**
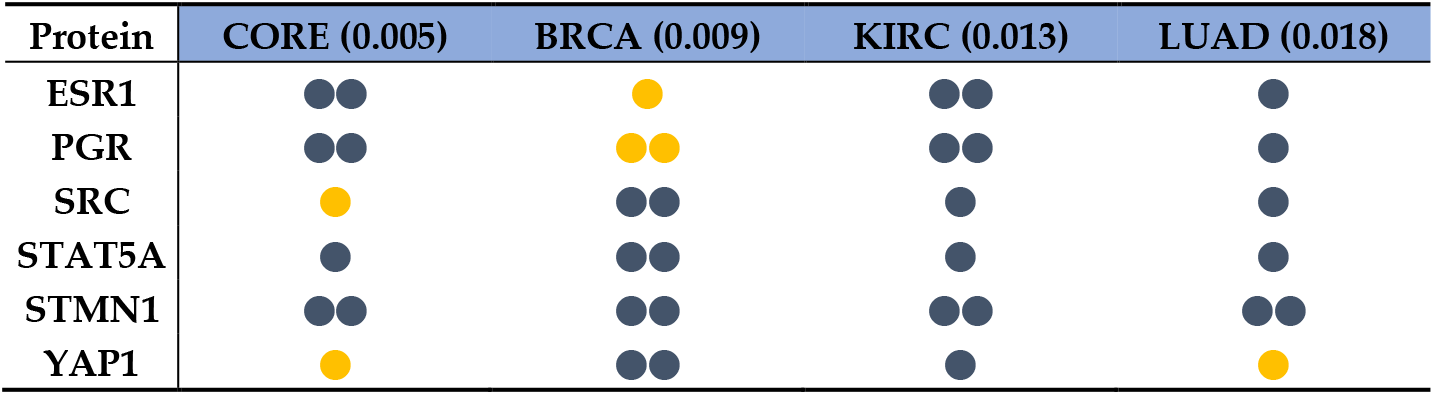
Differential protein expression in CORE, BRCA, KIRC, and LUAD for ERBB4-mediated nuclear signaling genes. Two blue or yellow dots indicate significantly higher protein expression in Blacks or Whites, respectively. A color-coded single dot represents non-significantly higher expression in the corresponding race.

In LUAD, TP53-mediated regulation of apoptotic gene transcription was increased in Black patients compared to Whites (adjusted p = 0.002): Blacks exhibited higher protein levels of ATM and BID, while TP53 and BAX expression was not significantly different between Blacks and Whites. In THCA, TP53 regulation of apoptotic gene transcription was increased in Asians compared to Whites (adjusted p = 0.017): Asians exhibited higher levels of BID protein. ATM, TP53, and BAX expression was also higher in Asians, although these differences were not significant.

In STAD, FOXO-mediated transcription of cell cycle genes was down-regulated in Asian patients compared to Whites (adjusted p = 0.014).

## 4. Discussion

In this study, we dissected large genomic, transcriptomic, proteomic, and survival datasets available from The Cancer Genome Atlas (TCGA) to identify molecular differences between White, Black, and Asian patients that might contribute to cancer health disparities. These findings help illustrate the complex nature of the tumor molecular differences that exist between racial groups, particularly between Whites and Blacks/African Americans. Importantly, there were insufficient data for extensive comparisons between Whites and Asians, highlighting the need for more equitable racial representation. Nonetheless, our methods provide a valuable pipeline to analyze the race dependency of DEGs, proteins, and pathways as well as their survival impact. We investigated the expression of 155 cancer-relevant genes and their negative or positive associations with patient survival to elucidate race-dependent molecular factors in multiple cancer types. We only included one dataset per cancer type (with the exception of breast cancer) to develop a proof of concept for the utility of such analyses that can be expanded to extract valuable information from even more cancer datasets.

Most clinically significant genes were either differentially expressed in one particular cancer type or not different between races. Even so, we identified several genes that were both differentially expressed and clinically significant in at least two cancer types. For example, the cell cycle progression gene *CCNE1* was both differentially expressed and implicated in patient survival in both breast and endometrial cancer. Additionally, the DNA repair gene MSH2 exhibited differential expression in kidney clear cell, liver, and endometrial cancer. *CCNB1, CCNE2*, and *FOXM1* were three other genes that were differentially expressed in more than one carcinoma: specifically, in kidney clear cell and liver carcinoma. The clinical implications of multiple members of the cyclin family of proteins as well as the cell cycle regulator *FOXM1* across multiple cancer types suggest that differences in cell cycle regulation are major contributors to the racial cancer disparity. Previous cancer-specific studies have also proposed this connection in breast [38] and endometrial carcinoma [24].

It is important to note that although we focused on genes that had a significant impact on overall survival, we also identified many other non-clinically significant genes that were differentially expressed between the examined races. It may be useful to perform further research on this subset of genes and their relevant pathways to uncover any unapparent relationships, if any, between expression and patient survival.

Additionally, the source datasets varied in their abundance of differentially expressed biomarkers. While several pathways were identified in multiple cancer types as being regulated in a race-dependent manner with direct correlation to altered molecular patterns, there was a near complete lack of molecular markers in others. For example, none of the clinically significant genes were differentially expressed between White and Asian patients in stomach adenocarcinoma.

A striking group of pathways that was significantly affected by race were DNA repair mechanisms. Most notably, racial differences between Black and White patients in the expression of genes pertaining to homology-directed repair, including *RAD50, RAD51, MRE11, ERCC1*, and *BRCA2*, was observed in five cancer types. Moreover, genes involved in nucleotide excision repair and DNA mismatch repair were differentially expressed in colorectal and liver cancer respectively. Nuclear signaling mediated by ERBB4 was upregulated in Blacks in four cancer types, a significant finding considering that overexpression of the kinase is associated with cancer development [39]. **Table 11** lists the major findings of this study, indicating the major pathways identified as differentially regulated.

**Table 11.**
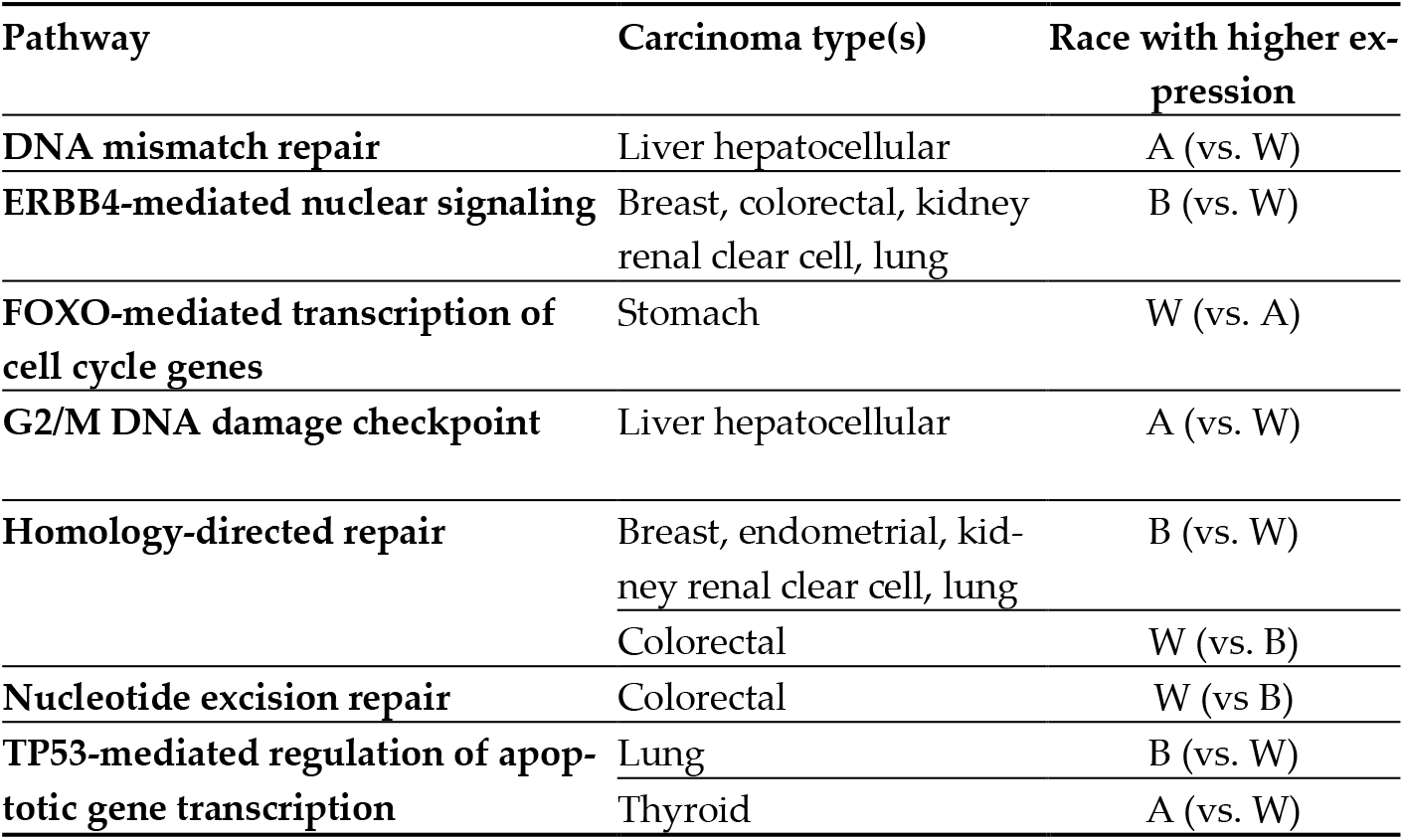
Differentially expressed pathways and their respective carcinoma types. A: Asian, B: Black, W: White.

| Pathway | Carcinoma type(s) | Race with higher expression |
| --- | --- | --- |
| DNA mismatch repair | Liver hepatocellular | A (vs. W) |
| ERBB4-mediated nuclear signaling | Breast, colorectal, kidney renal clear cell, lung | B (vs. W) |
| FOXO-mediated transcription of cell cycle genes | Stomach | W (vs. A) |
| G2/M DNA damage checkpoint | Liver hepatocellular | A (vs. W) |
| Homology-directed repair | Breast, endometrial, kidney renal clear cell, lung | B (vs. W) |
|  | Colorectal | W (vs. B) |
| Nucleotide excision repair | Colorectal | W (vs. B) |
| TP53-mediated regulation of apoptotic gene transcription | Lung | B (vs. W) |
|  | Thyroid | A (vs. W) |

Although protein data for many genes were absent from the original TCGA datasets, we can attempt to reconcile the mRNA differential expression analysis results with existing literature on protein imbalances across racial groups. In UCEC, *MDM2* mRNA expression was higher in White patients, a result consistent with the reported elevation of MDM2 protein expression in Caucasian Americans when compared to African Americans [40]. Additionally, increased *CDKN2A* transcript levels in Blacks correspond to the reported upregulation of CDKN2A in Black endometrial cancer patients. However, elevated transcript levels of *FGF12* in Blacks in UCEC conflict with the reported enrichment of fibroblast growth factor 12 in White endometrial cancer patients [24]. Such inconsistencies between mRNA and protein differential expression may be attributable to racial variation in post-transcriptional modifications, which may also play a role in the health disparity and is thus a promising topic for future research.

The autocrine-paracrine growth factor insulin-like growth factor 1 (IGF-1) has been reported to be a promoter of cancer that inhibits the sex-hormone binding globulin (SHBG) and exists in elevated levels in African American individuals [1]. Interestingly, although they were not identified as clinically significant for breast cancer, cBioPortal analysis revealed significantly decreased levels of *IGF1* mRNA and elevated levels of *SHBG* mRNA in Black patients in both BRCA-P and BRCA-C studies.

Although this study provides a methodological pipeline to identify clinically valuable molecular targets with preexisting datasets, we recognize certain limitations. The use of broadly defined racial groups and cancer types in our analyses neglects their great variability (e.g. different carcinoma sub-types.) There is also a consistent lack of representation of Black and Asian patients in the source datasets, an issue that must be adequately addressed by the broader research community. Additionally, in many cases, the transcriptome data does not accurately predict the corresponding proteomic data. The source data only included protein information for a few hundred genes, hence our pathway analyses are far from conclusive. We were not able to perform thorough analysis of the prostate adenocarcinoma dataset as it lacked survival data. Nevertheless, we highlight trends between gene survival impact and differential expression that suggest that disparities in cancer outcomes have direct molecular causes. Future research should work with larger tumor datasets with greater representation of Asian and Black/African American patients to yield more statistically accurate analyses. Ultimately, the cancer health disparity is a result of a combination of many factors including genetic polymorphisms, distinct molecular signatures, the synergistic effects of many traits, and lifestyle and environmental influences. With more equitable racial representation in tumor datasets and increased collection of proteomics data, we will be able to solidify our knowledge of molecular racial disparities and work towards the development of personalized cancer therapeutics. Genome projects such as the “1000 Genomes Project” [41] and the NIH-initiated “All of Us” [42] will provide more opportunities to identify unique molecular signatures in the future.

## 5. Conclusions

Racial patterns of differential gene expression and pathway regulation varied between different carcinomas. The expression of certain cell cycle genes (*CCNB1, CCNE1, CCNE2*, and *FOXM1*) were race-dependent in several cancer types and correlated with significant survival impacts. We also found racial variation in several DNA repair mechanisms and oncogenic pathways. Importantly, homology directed repair was upregulated in Black patients compared to White patients in breast, endometrial, kidney renal clear cell, and lung carcinomas. ERBB4-mediated nuclear signaling, which is associated with cancer development, was upregulated in Blacks in breast, colorectal, kidney renal clear cell, and lung carcinomas.

## Supplementary Materials

The following supporting information can be downloaded at: S1: List of 155 analyzed genes. S2: Complete differential mRNA expression results. S3: Complete differential protein expression results. S4: Complete differential methylation results. S5: Original Reactome report.

## Author Contributions

This paper was conceptualized, written, and reviewed by A.S. and B.L. B.L. designed the methodology, performed all data analyses, and prepared the original draft. A.S. provided overall supervision and guidance during data collection, analysis, and writing. X. J. reviewed and edited.

## Data Availability Statement

Data used in this article is publicly available and has been referenced. Computer code can be found at https://github.com/brian-lei/cancerracial-disparity.

## Acknowledgments

We thank Dr. Gianni Nigita (Department of Cancer Biology and Genetics and Comprehensive Cancer Center, The Ohio State University, Columbus, OH 43210, USA) for his valuable comments and insights on this study during the editing of this manuscript.

